## Supplementary info: technical notes for "Efficient and robust coding in heterogeneous recurrent networks"

### Supplementary Note 1: Technicalities

**A. A word on notation.** Firstly, we use the terms 'filter' and 'kernel' interchangably in this paper. Secondly, following the notation of amongst others Dayan and Abbott (89), we describe a spike-triggered average (STA) or an input filter as a filter where positive time $t$ denotes the time before the spike:

$$\text{STA}(t) = s(-t) * \rho(t) = \frac{1}{n}\sum_{i=1}^{n} s(t^i - t) \tag{18}$$

where the symbol $*$ stands for a convolution, $s(t)$ denotes the input signal and $t^i$ are the spike times of the spike train $\rho(t)$:

$$\rho(t) = \sum_{i=1}^{n} \delta(t - t^i)$$

**Solution 1: choose $\Delta$ smaller than the size of the filter** If the delay $\Delta$ is larger than the size of the filter, the problem explained above does not occur. This can be understood as follows: the MSE is calculated up to $T+\Delta$. However, if $\Delta$ exceeds the time of the filter $g$, there is no estimate between $T$ and $T+\Delta$, so any non-zero signal will add to the MSE. The system tries to reduce this error by firing more spikes, which results in an overestimation of the signal.

**Solution 2: gain modulation** A simple solution to the problem posed before, is increasing the threshold of the system with the parameter $\nu$. If we take $\nu$ to be slightly smaller than the average surface of the input filter

$$\nu = \frac{1}{2}\int_0^{T+\Delta} g_m(t-T)^2 dt - \epsilon, \tag{20}$$

the neuron will now spike at the right time. However, this also makes the neuron spike more sparsely and hence make a worse estimation of signals with a small amplitude. The threshold will have to be put even higher for larger-amplitude signals. Therefore, the threshold parameter $\nu$ should depend on the shape and amplitude of the input signal (ideally on the amplitude of the input signal convolved with the input filter). This assumes a mechanism of gain modulation that we would have to add to the model.

**Solution 3: adjusting the spike rule** A third solution is adjusting the spike rule, so that the system waits with firing a spike to see if the error is reduced more one time step later:

$$\Delta E = E^{\text{no spike at } T}(T+\Delta) - E^{\text{spike at } T}(T+\Delta) > 0 \wedge \frac{d\Delta E}{dt} <= 0 \tag{21}$$

which reduces to

$$V > \Theta + \nu \wedge \frac{dV}{dt} <= 0 \tag{22}$$

This spike rule can induce a bursting response to a step input. The system is now capable of responding optimally to a variety of inputs without the need to adjust a parameter line $\nu$. However, wIth this spike rule, the model is not GLM-like anymore. Moreover, the neuron cannot respond to slowly increasing inputs such as ramps.

**Solution 4: make an assumption about the autocorrelation of the input** The overestimation-problem arises, because at time $T+\Delta$, we only know the spikes up to $T$:

$$\begin{aligned} &\int_{t=0}^{T+\Delta} dt \left( g_k(t-T) \sum_{n=1}^{N} \sum_{t_n^i \neq T} g_n(t-t_n^i) \right) = \\ &\int_{t=0}^{T+\Delta} dt \left( \sum_{n=1}^{N} g_k(t-T) \sum_{t_n^i < t_k^l} g_n(t-t_n^i) \right) + \int_{t=0}^{t_k^l+\Delta} dt \left( \sum_{n=1}^{N} g_k(t-T) \sum_{t_n^i > T} g_n(t-t_n^i) \right). \end{aligned} \tag{23}$$

We can make an estimation for the second term on the right of equation (23). For instance, we can assume that the estimate changes slowly relative to the length of the filter, i.e. that the spike train of the network in $T < t < T+\Delta$ is the spike train of $T-\Delta < t < T$ mirrored. However, $g_k(t-T)g_n(t-t_n^i)$ is typically not symmetric in $t_n^i = T$. If we average over the surface of $g_k(t-T)g_n(t-t_n^i)$ before $t_n^i = T$ to compensate for this, this results in an output filter defined by

$$g_k^{\mathrm{o}} = (1+f_{kk})g_k^{\mathrm{i}} * g_k \tag{24}$$

and the lateral filters received by neuron k from neuron n defined by

$$g_{kn}^{\mathrm{l}} = (1+f_{kn})g_k^{\mathrm{i}} * g_n, \tag{25}$$

where

$$f_{kn} = \frac{\int_{\tau=0}^{T}\int_{t=0}^{T+\Delta} |g_k(t-T)g_n(t-\tau)| dt d\tau}{\int_{\tau=T}^{T+\Delta}\int_{t=0}^{T+\Delta} |g_k(t-T)g_n(t-\tau)| dt d\tau} \tag{26}$$

***B.2. Conclusion.*** The model can overestimate the signal, if the amount of neurons is very small and the delay $\Delta$ is the same size or larger than the nonzero part of the filter. Therefore, in this paper, we only used values of $\Delta$ that are smaller than the size of the filter. Alternatively, the threshold parameter $\nu$ can be adjusted to match the amplitude of the input. The alternative spike rule (22) does not need an adjustment of the threshold, but a neuron using this spike rule does not respond to slowly increasing input, or can respond too late to other inputs. Finally, assumptions about how the input changes can be made.
